# Whole brain wiring diagram of oxytocin system in adult mice

**DOI:** 10.1101/2020.10.01.320978

**Authors:** Seoyoung Son, Steffy B. Manjila, Kyra T. Newmaster, Yuan-ting Wu, Daniel J. Vanselow, Matt Ciarletta, Todd E. Anthony, Keith C. Cheng, Yongsoo Kim

## Abstract

Oxytocin (OT) neurons regulate diverse physiological responses via direct connections with different neural circuits. However, the lack of comprehensive input-output wiring diagrams of OT neurons and their quantitative relationship with OT receptor (OTR) expression presents challenges to understanding circuit specific OT functions. Here, we establish a whole-brain distribution and anatomical connectivity map of OT neurons, and their relationship with OTR expression using cell type specific viral tools and high-resolution 3D mapping methods. We utilize a flatmap to describe OT neuronal expression in four hypothalamic domains including under-characterized OT neurons in the tuberal nucleus. OT neurons in the paraventricular hypothalamus (PVH) broadly project to nine functional circuits that control cognition, brain state, and somatic visceral response. In contrast, OT neurons in the supraoptic (SO) and accessory nuclei have limited central projection to a small subset of the nine circuits. Surprisingly, quantitative comparison between OT output and OTR expression showed no significant correlation across the whole brain, suggesting abundant indirect OT signaling in OTR expressing areas. Unlike output, OT neurons in both the PVH and SO receive similar mono-synaptic inputs from a subset of the nine circuits mainly in the thalamic, hypothalamic, and cerebral nuclei areas. Our results suggest that PVH-OT neurons serve as a central modulator to integrate external and internal information via largely reciprocal connection with the nine circuits while the SO-OT neurons act mainly as unidirectional OT hormonal output. In summary, our OT wiring diagram provides anatomical insights about distinct behavioral functions of OT signaling in the brain.

**Significance Statement:** Oxytocin (OT) neurons regulate diverse physiological functions from pro-social behavior to pain sensation via central projection in the brain. Thus, understanding detailed anatomical connectivity of OT neurons can provide insight on circuit specific roles of OT signaling in regulating different physiological functions. Here, we utilize high resolution mapping methods to describe the 3D distribution, mono-synaptic input and long-range output of OT neurons, and their relationship with OT receptor (OTR) expression across the entire mouse brain. We found OT connections with nine functional circuits controlling cognition, brain state, and somatic visceral response. Furthermore, we identified a quantitatively unmatched OT-OTR relationship, suggesting broad indirect OT signaling. Together, our comprehensive OT wiring diagram advances our understanding of circuit specific roles of OT neurons.

## Introduction

Oxytocin (OT) is a highly conserved neuropeptide, playing key roles in regulating social behavior and other physiological functions (Althammer et al., 2018; Jurek and Neumann, 2018; Quintana and Guastella, 2020). Impairment in OT signaling has been heavily implicated in many neurodevelopmental disorders including autism (Francis et al., 2014; Rajamani et al., 2018). Altering OT signaling is being pursued as a potential therapy to alleviate social behavioral deficits in many brain disorders (Guastella and Hickie, 2015). However, our limited neural circuit based understanding of OT signaling in the brain hampers the development of targeted therapeutic approaches aimed at altering specific OT functions without affecting other biological pathways. A comprehensive anatomical understanding of OT neurons would enable integrated neural circuit specific studies to decipher the neural substrate of distinct OT functions.

The majority of OT producing neurons are located in the paraventricular nucleus of the hypothalamus (PVH) and the supraoptic nucleus (SO) while fewer OT neurons reside in the extended amygdala (Biag et al., 2012; Madrigal and Jurado, 2021). OT neurons receive input from distinct brain regions (e.g., the thalamus) and integrate sensory input with internal information to release OT in a context dependent manner in order to modulate specific downstream circuitry (Grinevich and Neumann, 2020; Tang et al., 2020). The actions of OT are mainly mediated by a single subtype of the OT receptor (OTR) expressed in distinct brain regions as well as peripheral tissues (Gimpl and Fahrenholz, 2001; Grinevich et al., 2016; Newmaster et al., 2020). In addition to the well-known peripheral release of OT as a hormone via the posterior pituitary, OT neurons send direct projections to specific brain areas that frequently express OTR, thereby modulating circuit specific functions (Grinevich et al., 2016; Liao et al., 2020). For example, OT signaling is linked with the medial prefrontal cortex for social cognition (Sabihi et al., 2014; Li et al., 2016), CA2 of the hippocampus for social memory (Raam et al., 2017; Tirko et al., 2018), the central amygdala for fear modulations (Knobloch et al., 2012; Ferretti et al., 2019), the parabrachial nucleus (PB) for fluid intake (Ryan et al., 2017), and the spinal cord for pain perception (Eliava et al., 2016; Boll et al., 2018). Despite these prior studies, we still lack a quantitative and comprehensive wiring diagram of the OT neurons in a standard 3D reference brain.

Here, we establish a comprehensive wiring diagram of OT neurons in the mouse brain using a high-resolution quantitative brain mapping method in combination with cell type specific transgenic mice and viral tools. All whole brain datasets are registered in the Allen Common Coordinate Framework (CCF) to facilitate data cross-comparison (Wang et al., 2020), and high resolution images can be easily viewed using a new web visualization (https://kimlab.io/brain-map/ot_wiring/). Using the new resource, we identified distinct OT neuronal connection with nine circuits that can explain diverse OT functions. Moreover, we found lack of quantitative correlation between OT output and OTR expression across the whole brain, suggesting abundant indirect OT signaling in OTR expressing brain areas.

## Material and Method

### Animals

All animal care and experimental procedures are approved by the Penn State University Institutional Animal Care Use Committee (IACUC). *Ot-Cre* mice (Choe et al., 2015) were originally produced in the Gloria B. Choi lab at the Massachusetts Institute of Technology and imported to the Penn State University (Kim Lab). To generate OT-Cre;Ai14 mice, *Ot-Cre* mice were crossed with Ai14 mice, expressing tdTomato following Cre-mediated recombination (Jax: 007914, C57Bl/6 J background). 2 months old C57Bl/6 J mice were used for whole brain tissue clearing and immunostaining. Mice received food and water ad libitum and were housed under constant temperature and light conditions (12 hrs light and 12 hrs dark cycle).

### Experimental design and statistical analyses

For OT neuron distribution mapping (Fig.1), we used 3 males, 3 females (virgin), and 2 females (lactating) of 2 – 4 month old OT-Cre;Ai14 mice with STPT imaging. We also used 4 males, 3 females (virgin) of 2 month old C57bl/6 mice for tissue clearing and LSFM imaging based quantification (Fig.1). Since we did not observe significant difference in OT neuronal number, we combined data from both sexes to generate representative cell counting (Table 1). For anterograde projectome mapping in 2 – 4 month old OT-Cre (Fig.2), we used 2 males, 3 females (virgin), and 3 females (lactating) with 500nl of AAV injection, and 3 males, 3 females (virgin), and 1 female (lactating) with 50-150nl of AAV injection for the PVH targeting. Moreover, we used 5 males and 5 females (virgin) for the SO, 3 males and 2 females (virgin) for the TU, 1 male and 1 female (virgin) for the AN. For oxytocin receptor expression mapping using OTR-Venus mice (Fig.3), we used 6 males and 8 females (virgin). For rabies input mapping (Fig.4), we used 3 males and 3 females (virgin) for the PVH, and 3 males and 1 female (virgin) for the SO.

**Figure 1.**
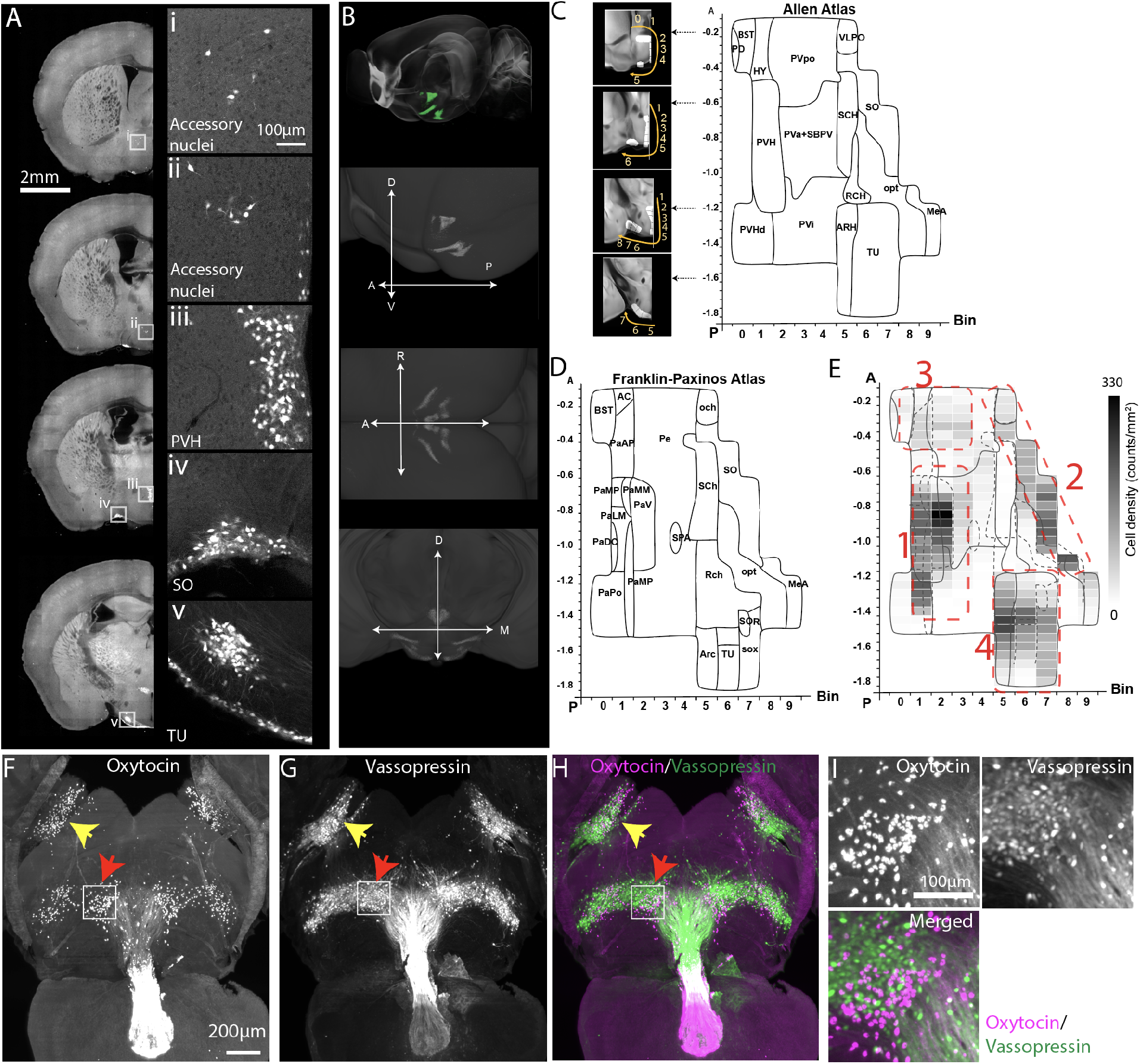
Distribution of oxytocin neurons. (A) Signals from OT-Cre;Ai14 mice across representative coronal planes of the mouse brain. Figures on the right column are high magnification images from white boxed areas in the left column. (B) (top) STPT images registered onto the Allen CCF reference brain. Green signals represent averaged OT neurons from 8 animals. (bottom) 3D distribution of oxytocin neurons. A/P: anterior/posterior, L/M: Lateral/medial, D/V: dorsal/ventral. See also Movie S1, Table 1. (C-D) 2D hypothalamic flatmaps. Small inserts with coronal sections illustrate how bins (while areas with numbers) were generated at different coronal planes. Anatomical labels in the flatmap are delineated based on Allen mouse brain atlas (C) and Franklin-Paxinos atlas (D). The X-axis is for bin numbers and Y-axis is for the bregma A/P axis. The full name of abbreviations can be found in Table 1. (E) Heatmap of oxytocin neuronal density in four clusters with the overlay of Allen and Franklin-Paxinos labels in solid and dotted lines, respectively. Red dotted lines for four OT expressing domains. 1: PVH, 2: SO, 3: AN, and 4: TU. (F-I) Light sheet fluorescence microscopy imaging of whole brain immunostaining with OT and vasopressin antibodies. 500 μm thick z maximum projection of OT (F), vasopressin (G), and both (H). Yellow and red arrows for the SO and the TU, respectively. (I) high magnification images from the white boxed TU area in (F-G), Note the lack of colocalization between the OT and vasopressin.

**Figure 2.**
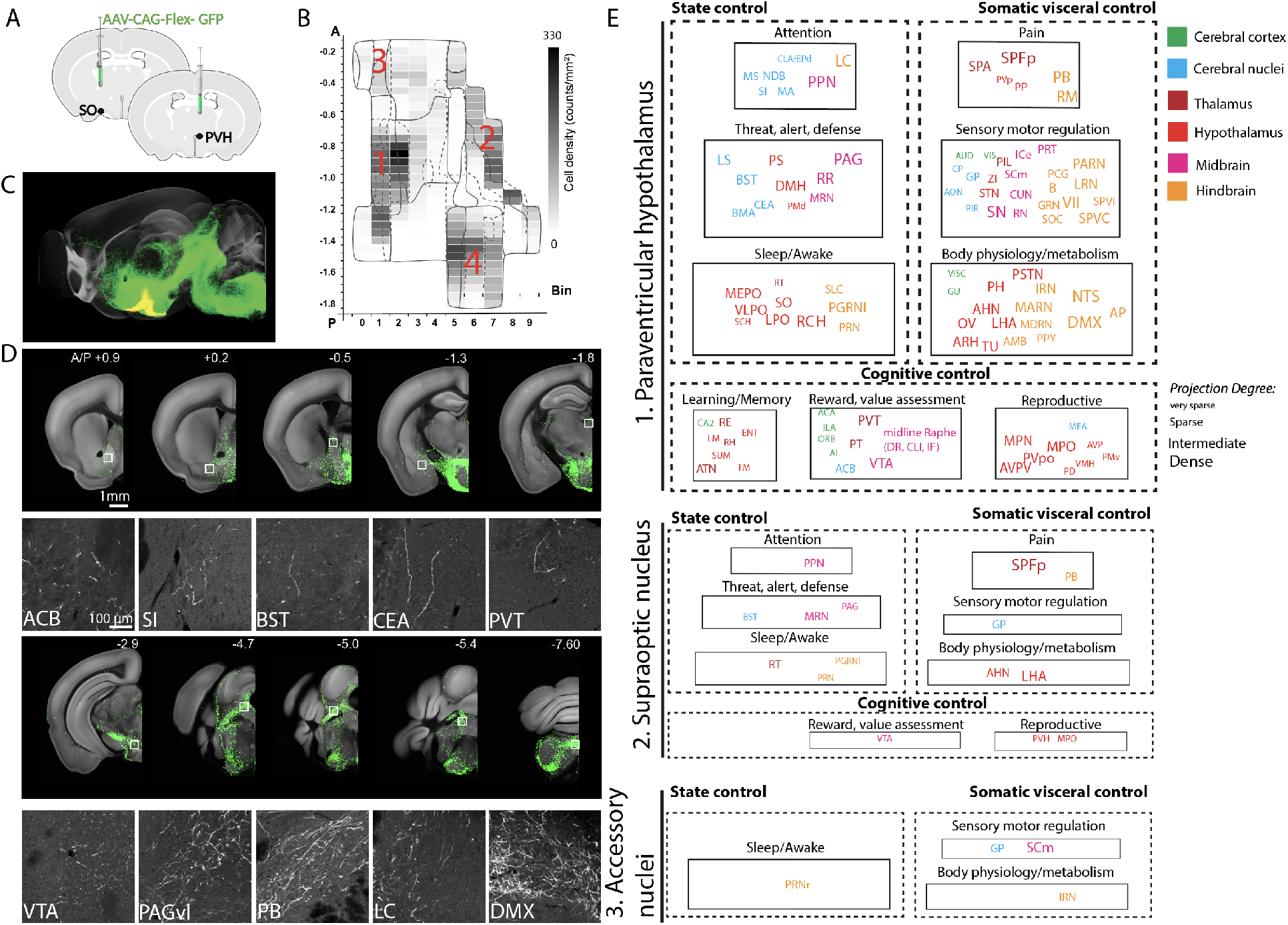
Anterograde projection map of oxytocin neurons. (A) Conditional AAV-GFP was injected in OT neuron containing hypothalamic areas. (B) Four major areas of viral injections, 1: the PVH, 2: the SO, 3: the AN, 4: the TU area. (C) Projection outputs from the PVH (green) and SO (yellow) oxytocin neurons registered in the Allen CCF. See also Movie S2. (D) Examples of long-range projections (green) from OT neurons in the PVH. The bottom panel is high mag images from white boxed areas in the top panel. (E) Nine functional circuits that receive long-range projection from OT neurons in the three different injection area 1 - 3. Color and size of each region of interest represent anatomical ontology and the abundance (degree) of the projection. The full name of abbreviations can be found in Table 2.

**Figure 3.**
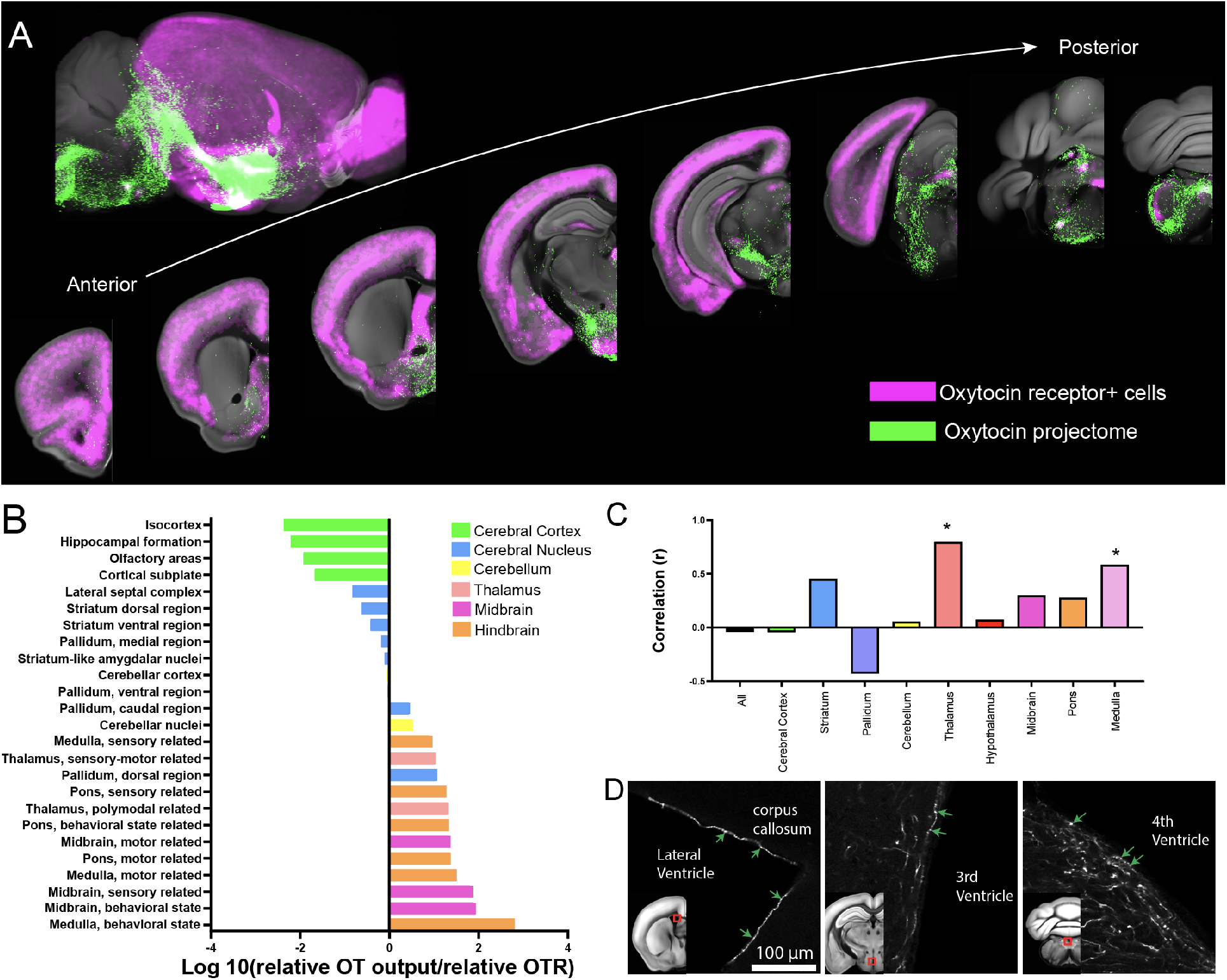
Comparison between the oxytocin output and oxytocin receptor expression. (A) Composite images of representative OT neuronal projection outputs (red: combined from both the PVH and the SO) and OTR expression (green) across the mouse brain. See also Movie S3. (B) Quantitative comparison of relative OT projection pattern and OTR expression. Note that the cerebral cortex has very small OT/OTR ratio while the hindbrain and the midbrain shows higher ratio. (C) Correlation between OT projection and oxytocin receptor density (Spearman nonparametric correlation, *: p<0.05). Note no significant correlation across the whole brain areas despite the significant correlation in the thalamus and the medulla. (D) Examples of OT long range projection touching the surface of all major ventricles.

**Figure 4.**
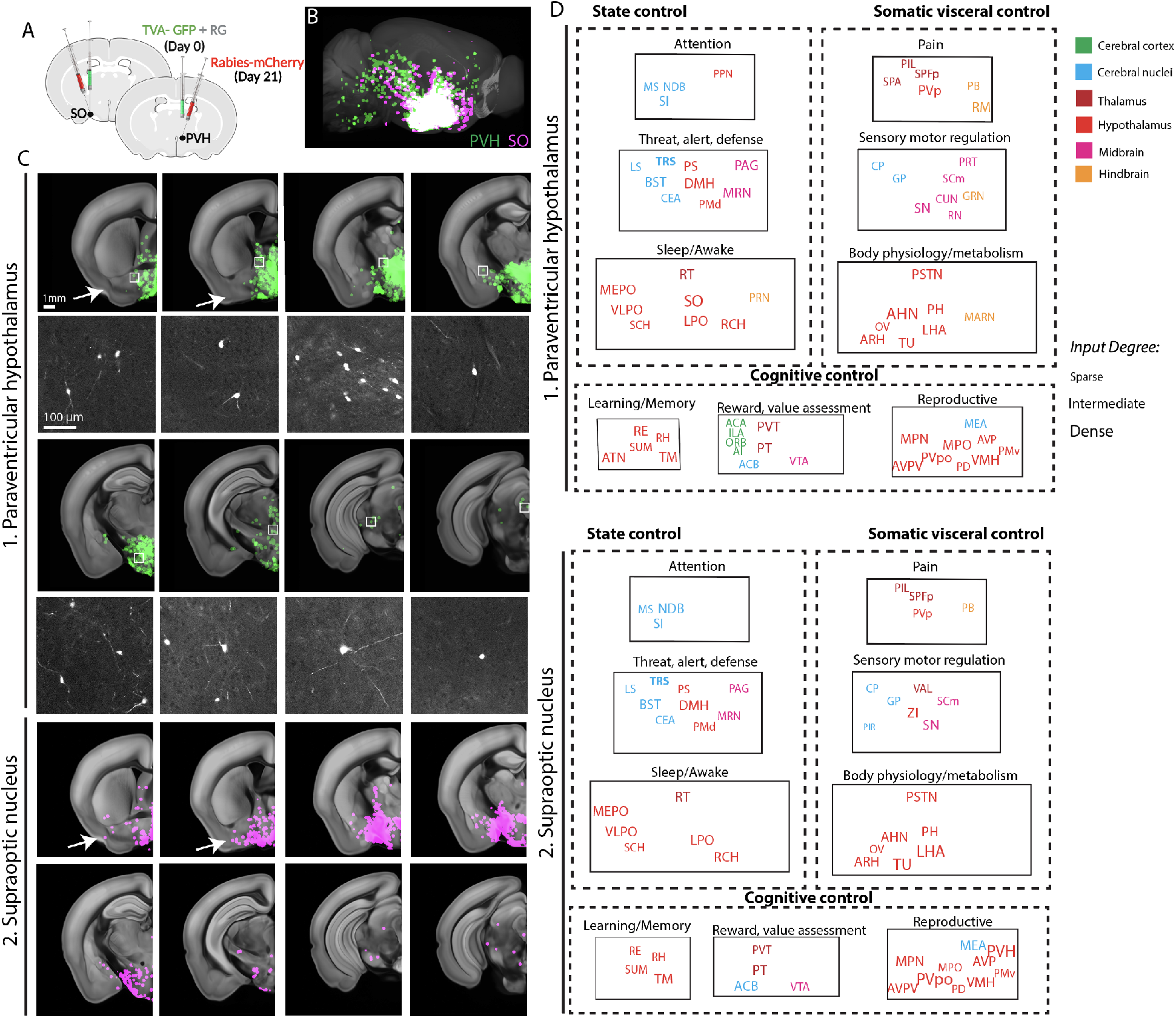
Monosynaptic input map of oxytocin neurons in the PVH and the SO. (A) Conditional mono-synaptic tracing rabies virus was injected in the PVH or the SO of the OT-Cre mice. (B) Brain-wide inputs into the PVH (green, n = 6 animals) and SO (red, n = 4 animals) OT neurons. The maximum signals of all samples from each anatomical region were overlaid on the reference brain. See also Movie S4. (C) Representative mono-synaptic inputs in different coronal planes (top panel) and high mag images from white boxed areas (bottom panel). Arrows highlight input from lateral brain areas for the SO compared to the PVH. (D) Nine functional circuits that provide mono-synaptic input to OT neurons in the two anatomical areas. Note overall similar input patterns for both areas. The full name of abbreviations can be found in Table 2.

**Table 1.** Oxytocin neuron counting. Transgenic animal counting is from serial two-photon tomography imaging of OT-Cre;Ai14 mice (n=8) and 3D immunolabeling counting is from light sheet fluorescence microscopy imaging of C57 after tissue clearing and 3D oxytocin antibody staining (n=7). Counting data are mean ± standard deviation.

| Cluster | Full Names | Abbreviations | Transgenic | 3D immuno labeling |
| --- | --- | --- | --- | --- |
| **1** | Paraventricular hypothalamic nucleus | PVH | 511.5 ± 147.1 | 818 ± 169 |
|  | Periventricular hypothalamic nucleus, anterior part | PVa | 47.6 ± 22.7 | 119 ± 23 |
|  | Periventricular hypothalamic nucleus, intermediate part | PVi | 9.6 ± 11.4 | 14 ± 5 |
|  | Subparaventricular zone | SBPV | 17.9 ± 20.8 | 113 ± 49 |
|  | Paraventricular hypothalamic nucleus, descending division | PVHd | 153.8 ± 49.7 | 27 ± 16 |
| **2** | Supraoptic nucleus | SO | 202.3 ± 65.5 | 654 ± 89 |
|  | Medial amygdalar nucleus | MEA | 108.7 ± 49.6 | 10 ± 4 |
|  | Ventrolateral preoptic nucleus | VLPO | 21.9 ± 23.3 | 50 ± 20 |
| **3** | Bed nuclei of the stria terminalis | BST | 27.8 ± 11 | 68 ± 34 |
|  | Periventricular hypothalamic nucleus, preoptic part | PVpo | 51.1 ± 18.3 | 194 ± 40 |
|  | Substantia innominata | SI | 6 ± 3.1 | 2 ± 2 |
|  | Medial preoptic nucleus | MPN | 18 ± 5.8 | 71 ± 14 |
|  | Lateral hypothalamic area | LHA | 45.9 ± 14.4 | 32 ± 12 |
|  | Lateral preoptic area | LPO | 2.1 ± 2 | 4 ± 2 |
| **4** | Tuberal nucleus | TU | 472.9 ± 65.2 | 393 ± 121 |
|  | Arcuate hypothalamic nucleus | ARH | 147.5 ± 86.5 | 272 ± 83 |

**Table 2.** Nine functional circuits that are connected with oxytocin neurons.

| Functional Circuits | Full Name | Abbreviation |
| --- | --- | --- |
| **Attention** | Medial septal nucleus | MS |
|  | Diagonal band nucleus | NDB |
|  | Substantia innominata | SI |
|  | Medial septal nucleus | MS |
|  | Clastrum/dorsal endopiriform nucleus | CLA/EPd |
|  | Locus ceruleus | LC |
|  | Pedunculopontine nucleus | PPN |
| **Threat, alert, defense** | Lateral septal nucleus | LS |
|  | Bed nuclei of the stria terminalis | BST |
|  | Central amygdalar nucleus | CEA |
|  | Basal medially amygdala | BMA |
|  | Parastrial nucleus | PS |
|  | Dorsomedial nucleus of the hypothalamus | DMH |
|  | Dorsal premammillary nucleus | PMd |
|  | Periaqueductal gray, ventral lateral | PAG |
|  | Midbrain reticular nucleus, retrorubral area | RR |
|  | Midbrain reticular nucleus | MRN |
| **Sleep/Awake** | Median preoptic nucleus | MEPO |
|  | Ventrolateral preoptic nucleus | VLPO |
|  | Suprachiasmatic nucleus | SCH |
|  | Reticular Thalamus | RT |
|  | Supraoptic nucleus | SO |
|  | Lateral preoptic area | LPO |
|  | Subceruleus nucleus | SLC |
|  | Retrochiasmatic area | RCH |
|  | Paragigantocellular reticular nucleus, lateral part | PGRNI |
|  | Pontine reticular nucleus | PRN |
| **Pain** | Subparafascicular area | SPA |
|  | Subparafascicular nucleus, posterior | SPFp |
|  | Periventricular hypothalamic nucleus, posterior part | PVp |
|  | Peripeduncular nucleus | PP |
|  | Parabrachial nucleus | PB |
|  | Nucleus raphè magnus | RM |
| **Sensory motor regulation** | Auditory cortex | AUD |
|  | Visual cortex | VIS |
|  | Caudate Putamen | CP |
|  | Globus pallidus | GP |
|  | Anterior olfactory nucleus | AON |
|  | Piriform cortex | PIR |
|  | Posterior Intralaminar Thalamic nucleus | PIL |
|  | Zona Incerta | ZI |
|  | Subthalamic nucleus | STN |
|  | Substantia nigra | SN |
|  | Inferior colliculus, external nucleus | ICe |
|  | Superior colliculus, motor | SCm |
|  | Cuneiform nucleus | CUN |
|  | Red nucleus | RN |
|  | Pretectal region | PRT |
|  | Pontine central gray | PCG |
|  | Barrington's nucleus | B |
|  | gigantocellular reticular nucleus | GRN |
|  | Superior olivary complex | SOC |
|  | Parvocellular reticular nucleus | PARN |
|  | Lateral reticular nucleus | LRN |
|  | Facial motor nucleus | VII |
|  | Spinal nucleus of the trigeminal, caudal part | SPVC |
|  | Spinal nucleus of the trigeminal, interpolar part | SPVI |
| **Body**<br>**physiology/metabolism** | Visceral cortex | VISC |
|  | Gustatory cortex | GU |
|  | Parasubthalamic nucleus | PSTN |
|  | Posterior hypothalamic nucleus | PH |
|  | Anterior hypothalamic nucleus | AHN |
|  | Vascular organ of the lamina terminalis | OV |
|  | arcuate hypothalamic nucleus | ARH |
|  | Lateral hypothalamic area | LHA |
|  | Tuberal nucleus | TU |
|  | Intermediate reticular nucleus | IRN |
|  | Magnocellular reticular nucleus | MARN |
|  | Medullary reticular nucleus | MDRN |
|  | Nucleus ambiguus | AMB |
|  | Parapyramidal nucleus | PPY |
|  | Nucleus of the solitary tract | NTS |
|  | Dorsal motor nucleus of the vagus nerve | DMX |
|  | Area postrema | AP |
| **Learning and Memory** | Field CA2 | CA2 |
|  | Nucleus of reunions | RE |
|  | Lateral mammillary nucleus | LM |
|  | Rhomboid nucleus | RH |
|  | Supramammillary nucleus | SUM |
|  | Anterior group of dorsal thalamus | ATN |
|  | Entorhinal area | ENT |
|  | Tuberomammillary nucleus | TM |
| **Reward, value**<br>**assessment** | Anterior cingulate area | ACA |
|  | Infralimbic cortex | ILA |
|  | Orbital cortex | ORB |
|  | Agranular insular area | AI |
|  | Nucleus accumbens | ACB |
|  | Paraventricular nucleus of the thalamus | PVT |
|  | Parataenial nucleus | PT |
|  | Dorsal Raphe | DR |
|  | Central linear nucleus raphe | CLI |
|  | Interfascicular nucleus raphe | IF |
|  | Ventral tegmental area | VTA |
| **Reproductive** | Medial preoptic nucleus | MPN |
|  | Periventricular hypothalamic nucleus, preoptic part | PVpo |
|  | Anteroventral periventricular nucleus | AVPV |
|  | Medial preoptic area | MPO |
|  | Anteroventral preoptic nucleus | AVP |
|  | Ventral premammillary nucleus | PMv |
|  | Ventromedial hypothalamic nucleus | VMH |
|  | Posterodorsal preoptic nucleus | PD |
|  | Medial amygdala | MEA |

To determine the correlation between OT area normalized projection and OTR density (Fig. 3C), we first tested for the normality of the data using the D’Agostino-Pearson normality test. Based on the normality test result, we performed Spearman nonparametric correlation test. GraphPad Prism 8 was used for all statistical analysis and graphs.

### Stereotaxic surgery and virus injections

*Ot-Cre* mice (8-11 weeks old, males and females) were anesthetized with isoflurane (controlled with Somnosuite, Kent Scientific) and mounted on a stereotaxic instrument (Angle Two, Leica) with a heating pad placed underneath. All injections were performed with pulled micropipettes (VWR, Cat# 53432-706). Through the small opening of the micropipette, virus was delivered at a rate of 75-100 nL per minute. The speed and volume of injection were monitored along with the calibration marks on the micropipette (1 mm =100 nL). To target the PVH, coordinates are anteroposterior (AP) from the Bregma: −0.58 mm; mediolateral (ML): 0.27 mm; dorsoventral (DV): −4.75 mm. Anterior PVH and posterior PVH injection coordinates are −0.35 mm (AP), 0.3 mm (ML), and −4.5 mm (DV) and −0.94 mm (AP), 0.39 mm (ML), and −4.55 mm (DV), respectively. Coordinates for the SO are −0.66 mm (AP), 1.3 mm (ML), and −5.8 mm (DV). For anterograde tracing, 50-500 nL of AAV2-CAG-Flex-EGFP virus (titer 3.7 × 10^12^ vg/ml, purchased from UNC vector core) was injected into the PVH (500 nL for maximum coverage, 50-150 nL for PVH subregion) and 150 nL of the virus was injected into the SO. Mice were euthanized three weeks later with Ketamine (100 mg/kg) and Xylazine (10 mg/kg) mixture. For monosynaptic retrograde labeling, 50-500 nL of rAAV1-synp-DIO-STPEPB (titer 3.9 × 10^12^ cg/ml, purchased from UNC vector core, a gift from Ian Wickersham (Addgene plasmid # 52473; http://n2t.net/addgene:52473; RRID:Addgene_52473 (Kohara et al., 2014)) was injected into the PVH, followed by the same quantity of EnvA G-deleted Rabies-mcherry virus (titer: 8.12×10^8^ TU /ml, purchased from the Salk Institute Viral Vector Core, a gift from Edward Callaway (Osakada et al., 2011), RRID:Addgene_32636) three weeks later into the same location. Mice were euthanized 7-8 days later with Ketamine (100 mg/kg) and Xylazine (10 mg/kg) mixture.

Control experiments were performed by injecting 500nl of rAAV1/Synp-DIO-STPE (titer 4.3 × 10^12^ vg/ml, purchased from UNC vector core, a gift from Ian Wickersham (Addgene plasmid # 52474; http://n2t.net/addgene:52474; RRID:Addgene_52474) (Kohara et al., 2014)) in *Ot-Cre* mice at the same co-ordinates for PVH and SO (N=1 each). EnvA G-deleted Rabies-mcherry virus was injected 3 weeks later and the mice were euthanized 7 days later for brain collection.

To check for leakiness of TVA, 500nl of rAAV1-synp-DIO-STPEPB was injected along with 50ul of 1:4 diluted pAAV-CAG-tdTomato (codon diversified) (a gift from Edward Boyden (Addgene plasmid # 59462; http://n2t.net/addgene:59462; RRID:Addgene_59462) in C57 mice at the same co-ordinates for PVH.

For the optimized G and split TVA tracing, AAV5-CAG(del)>TCIT(-ATG)-Flex(*loxP*)-SV40 (1:8 dilution, titer: 1.6 E+12gc/ml, a gift from Todd Anthony at Harvard) was co-injected with AAV5-CAG(del)>nC2oG-Flex(loxP) (titer: 2.8 E+12gc/ml, a gift from Todd Anthony) in *Ot-Cre* mice (N=2 each for PVH and SO co-ordinates) (150nl per brain). After 14 days, the same volume of EnvA G-deleted Rabies-EGFP virus (titer:4.89E+09 TU/ml, purchased from Salk Institute viral vector core, a gift from Edward Callaway (Addgene plasmid # 32635; http://n2t.net/addgene:32635; RRID:Addgene_32635) (Osakada et al., 2011)) was injected into the same location. The mice were euthanized 7 days later for brain collection and STPT imaging.

### STPT imaging and related data analysis

Transgenic or virus injected mice were transcardially perfused with 4% paraformaldehyde (PFA) in 0.1M phosphate buffer (PB, pH 7.4) after 0.9% saline. Brains were dissected out and post-fixed in 4% PFA overnight at 4°C. Fixed brains were stored in 0.05 M phosphate buffer at 4°C until imaged. To image the entire brain, serial two-photon tomography (TissueCyte 1000; Tissuevision) was used as previously described (Ragan et al., 2012; Kim et al., 2017; Newmaster et al., 2020). Briefly, the brain was embedded in 4% oxidized agarose and cross-linked with 0.2% sodium borohydride solution. The brain was imaged as 12 × 16 × 280 tiles with 1 × 1 μm^2^ *x,y* pixel resolution in every 50 μm *z*-section. We used 910 nm wavelength for two-photon excitation to excite both green (e.g., eGFP) and red signals (e.g., tdTomato). Signals were separated with 560 nm dichroic mirror and two band path filters (607/70-25 for red and 520/35-25 for green). Imaging tiles in each channel were stitched with custom-built software (Kim et al., 2017; Newmaster et al., 2020).

For quantitative projection data analysis, we used our previously published pipeline (Jeong et al., 2016). Briefly, both signal and background channels were z-normalized. Then, the background channel images were subtracted from the signal channel images to increase signal-to-noise ratio. Then, projection signals were converted to a binary map by applying an optimized threshold (8x standard deviation) to detect signals while minimizing noise from background autofluorescence. Then, binarized signals in each pixel were counted in 20 × 20 (*x,y*) pixel unit (voxel) and the value was assigned the corresponding voxel across the brain, which is defined as “projection strength”. Thus, range of the projection strength in a given voxel is between 0 and 400. Projection strength of each area is calculated by summing up all projection strength within an anatomically defined area. Autofluorescence of brains was used to register each brain to the Allen CCF using Elastix (Klein et al., 2010), then, the projection signals were transformed to the reference brain. Then, we used maximum projection of registered long-range output datasets from each area to create a representative projection data for further quantitative analysis (Movie S2). “Area normalized projection” represents normalized occupancy of projection signals in the ROI by dividing the projection strength with a total number of voxels in each ROI. For example, if total voxel count for one ROI was 20,000 and our projection strength showed 2,000 in the ROI, it will be (2,000/20,000)*100 = 10%. Regions with a projection strength greater than 1% is designated as dense, between 1 and 0.5 as intermediate, between 0.5 and 0.1 as sparse, and less than 0.1 as very sparse (Fig. 2E).

For cell counting analysis, we used a machine-learning algorithm to detect fluorescently labeled cells (Kim et al., 2017; Newmaster et al., 2020). The cell density in 2D (count/mm^2^) was calculated by dividing cell number with ROI area. 2D counting numbers were also converted into 3D counting using our previously calculated 3D conversion factor (1.4 for tdTomato) (Kim et al., 2017). To measure the volume of anatomical ROI, the reference Allen CCF was reverse registered onto individual brains using the Elastix. “Cell density (counts/mm^3^)” was calculated by dividing detected cell numbers in 3D with the anatomical ROI volume. The cell counting analysis was applied to OT-Cre;Ai14 and OTR-Venus cell distribution and inputs to the OT neurons. We used an average of individual datasets to create representative OT (OT-Cre;Ai14, Movie S1) and OTR (OTR-Venus, Movie S3) distribution and maximum projections to create mono synaptic input for OT neurons (rabies). For rabies input degree (Fig. 4D, Movie S4), Regions more than 100 cells are designated as dense, between 100 and 10 as intermediate, and less than 10 as sparse.

To compare relative abundance between OT output and OTR expression in Fig. 3C., relative cell density or output data in each region was calculated by dividing each data by summed density or output data from all areas (excluding viral injection sites), respectively. Then, log10 (relative OT output/relative OTR) was used to examine the quantitative relationship between the two signals.

### 2D hypothalamic and PVH Flatmap

To generate the hypothalamic flatmap, we adapted the previously used method (Kim et al., 2017) and applied it to the hypothalamic region. First, we created a binary image in the hypothalamic area based on the oxytocin expression. Second, a zero line was placed to generate evenly spaced bins along the dorsal to the ventral direction of the PVH and laterally extended to include TU and MEA at different coronal plains. To capture signals on the flatmap, bins were registered into the reference brain and the cell number in each bin was quantified as described before in the STPT data analyses section. Lastly, the mean number of the OT neurons in 8 OT-Cre;Ai14 brains were plotted in each flatmap using a custom-built matlab code. For the PVH flatmap, we followed the same procedure to generate a hypothalamic flatmap except for bin generation. Instead of delineating bins in a binary image, we assigned bin numbers in the PVH subregion of Franklin-Paxinos atlas (Paxinos and Franklin, 2008) in the dorsal to the ventral direction.

### Whole brain clearing and immunostaining, light sheet microscopy, and cell counting

C57Bl/6 J mice (4 males and 3 females at P56) were transcardially perfused with 0.9% saline followed by 4% PFA in 0.1M phosphate buffer (PB, pH 7.4). The decapitated heads were postfixed in 4% PFA overnight at 4°C and brains were dissected out the following day. All the following steps were performed on an orbital shaker unless otherwise specified. Dissected brains were delipidated in SBiP buffer (0.2mM Na2HPO4, 0.08% SDS (Sodium Dodecyl Sulfate), 0.16% 2-methyl 2-butanol, 0.08% 2-propanol). Delipidation was performed with 3-4 washes (10ml per wash) in SBiP for 24 hrs followed by one 10ml wash with SBiP for the next 4 days.

Samples were then moved to B1n buffer (0.1%v/v Tritox-X-100, 1% wt/v glycine, 0.001N NaOH, 0.008% wt/v sodium azide) for 1 day (10ml) and then shifted to 37^0^C incubation for 3 hrs. Once delipidation was completed, the samples were washed in PTwH (Tween 20-2ml, 10mg/ml Heparin-1ml and sodium azide-2g, made to 1L with 0.1M phosphate buffered saline) 3-5 times at 37^0^C for 24 hrs. The samples were then incubated in antibody solution (5% DMSO and 3% Donkey serum in PTwH-4ml per sample) containing primary antibodies for OT (ImmunoStar Cat# 20068, RRID:AB_572258, 1:500) and Vasopressin (Peninsula Laboratories Cat#T-5048, RRID:AB_2313978, 1:1000) for 10 days at 37^0^C. Next, PTwH washes were performed 4-5 times for 24 hrs at 37^0^C, followed by secondary antibody incubation. Secondary antibodies (1:500) were used as follows: Alexa Fluor 594 AffiniPure Fab Fragment Donkey Anti-Rabbit IgG (H+L) (Cat # 711-587-003, RRID: AB_2340623) and Alexa Fluor 647 AffiniPure F(ab’)_2_ Fragment Donkey Anti-Guinea Pig IgG (H+L) (Cat # 706-606-148, RRID:AB_2340477) in antibody solution (4ml per sample) for 10 days at 37^0^C. The samples were further washed 3-4 times in PTwH for 24 hrs at 37^0^C. Once immunolabeling was completed, the samples were moved to room temperature (RT) and further processed for tissue clearing. All the following steps were performed in a fume hood in glass containers and the containers were filled completely. Samples were dehydrated in the following series of methanol dilutions: 20%v/v-1 hr at RT, 40%v/v-1 hr at RT, 60%v/v-1 hr at RT, 80%v/v-1 hr at RT, 100%v/v-1 hr at RT and 100%v/v at RT overnight. Next, the samples were incubated for 3 hrs in 66%v/v dichloromethane/33%v/v Methanol at RT followed by 100% dichloromethane (Sigma Cat# 270997) incubations of 30min and 2hrs. Samples were then index matched in benzyl ether (Sigma Cat#108014) overnight without shaking. Once the samples are completely transparent (1-2 days), samples were moved to ethyl cinnamate (Sigma Cat#112372). Whole brain samples were then imaged using a light sheet microscope (SmartSPIM, Life Canvas) at 4x resolution.

OT cell detection and 3D counting workflow are similar to the STPT based quantification by applying a 2-D FFT high pass filter, normalizing the data by dividing it by the filtered part, thresholding and 3D water-shedding to find the mask of each cell, and finally documenting each cell with its centroid location.

### Immunohistochemistry, microscopic image, and cell counting

For immunohistochemistry, fixed brains were either embedded in 3% agarose or frozen after sinking in 30% sucrose in 0.2 M Phosphate buffer. Embedded or frozen brains were then cut on a vibratome (Leica vt1000s) or a microtome (Leica SM2010 R) at 50 μm thickness. Sections were stored at −20°C in a cryoprotectant solution (30% sucrose and 30% glycerol in 0.1 M PB) until immunostaining. For oxytocin staining, sections were washed three times in 1x PBS. After 1 hour incubation in blocking solution (10% donkey serum and 0.1 % Triton X-100), slices were incubated with oxytocin primary antibody (ImmunoStar Cat# 20068, RRID:AB_572258, 1:1000) in blocking solution for overnight at 4 °C. Sections were then washed three times with 1x PBS and further incubated in secondary antibodies (Thermo Fisher Scientific Cat# A-21206, RRID:AB_2535792, 1:500) for 1 hour at room temperature. After washing three times, slices were mounted onto slides and coverslipped with vectashield mounting media (Vector laboratories, H-1500-10). For microscopic imaging, a BZ-X700 fluorescence microscope (Keyence) and a confocal microscope (Zeiss 510) were used. A low magnification objective lens (4x) was used to image with a large enough view to define brain anterior-posterior location from bregma and higher magnification objective lenses (10x ~ 40x) were used to image sections depending on the cell density. Images were delineated manually based on the Franklin-Paxinos atlas and fluorescently tagged cells were manually quantified using the cell counter plug-in in FIJI (ImageJ, NIH).

### Software Accessibility

All custom-built codes and flatmaps used in the current study will be freely available upon request and can be used without any restriction.

#### Data Sharing Plan

Data files for the anterograde projectome, rabies based monosynaptic input, and OTR expression data registered on the Allen CCF are included as supplementary data.

High-resolution serial two-photon tomography images will be deposited in BrainImageLibrary (https://www.brainimagelibrary.org/) and web visualization link will be added upon publication.

## Results

### Quantitative density mapping of oxytocin neurons reveals four clusters in the adult mouse brain

We first aim to determine quantitative brain-wide OT distribution in complex 3D structures. To examine the anatomical distribution of OT neurons across the whole brain, we used OT knock-in mice with Cre recombinase (*Ot-Cre*) crossed with Ai14 reporter mice (OT-Cre;Ai14-heterozygotes) (Choe et al., 2015). We imaged the entire mouse brain at cellular resolution using serial two-photon tomography (STPT) and performed quantitative mapping using previously established computational methods (n=8 brains, Fig. 1A-B, Movie S1) (Kim et al., 2017). The PVH regions (PVH, descending division of PVH, anterior, intermediate, and subparaventricular zone) contain the highest density of OT neurons (~39%, 742 out of total 1899 cells) followed by the tuberal nucleus (TU), SO, and other areas (Table 1). To further visualize the spatial expression pattern of OT neurons, we created a flatmap (Fig. 1C). Evenly spaced bins provide a flattened 2D spatial unit to quantify and to display signals from the 3D brain. The flatmap was delineated with Allen Common Coordinate Framework (Allen CCF) and Franklin-Paxinos atlas based anatomical labels (Fig. 1C-D) (Paxinos and Franklin, 2008; Chon et al., 2019; Wang et al., 2020). The regional boundaries of the two labeling systems generally agreed with each other in the major OT expressing regions (e.g., the PVH and the SO) despite noticeable discrepancies in the caudal hypothalamic area (e.g., the TU) (Fig. 1C-D) (Chon et al., 2019). The OT density heatmap on the hypothalamic flatmap clearly shows four clusters: 1. the PVH, 2. the SO, 3. accessory nuclei (AN), 4. the TU area (Fig. 1E) (Knobloch and Grinevich, 2014). Notably, the largely overlooked TU area contains almost as high density of OT neurons as the PVH area (Fig. 1E).

To distinguish neurons actively expressing OT in adults from developmentally labeled cells, we performed immunohistochemistry using an OT antibody in OT-Cre;Ai14 mice. We confirmed that almost all OT immuno positive neurons (97%, 1733 out of 1790 cells, n=4 animals) were labeled by tdTomato from OT-Cre;Ai14 mice (Fig. S1). In contrast, 76% of tdTomato labeled cells were OT immuno positive (1733 out of 2277 cells) in the PVH. Smaller portions of tdTomato cells in the SO (44%, 654 out of 1508 cells) and the MEA (8%, 31 out of 375 cells) retain active OT expression (Fig. S1). This result suggested that OT neurons undergo OT expression changes during neurodevelopmental processes (Madrigal and Jurado, 2021).

To cross validate active expression of OT in the adult brain, we performed tissue clearing followed by 3D immunolabeling with OT and vasopressin antibodies in 8 weeks old C57bl/6 mice (n= 7 brains; Fig. 1F-I) (Renier et al., 2016). We developed 3D counting and image-registration methods to achieve similar unbiased brain-wide cell counting as done with STPT imaging (see Methods for more details). We observed similar OT staining distribution and overall slightly higher counting compared to our transgenic based mapping results (Table 1). For example, the estimated number of OT neurons in the PVH with the immunostaining was 1,095 cells out of total 3149 cells (~35%), which is higher than our transgenic based estimate mostly likely due to sensitive labeling based on antibody detection. Importantly, we confirmed the robust OT expression in the TU area that was not colocalized with vasopressin staining (Fig. 1I).

### Quantitative whole brain projection mapping of OT neurons reveals broad long-range projections in nine functional circuits

Next, we aim to establish a comprehensive anterograde projection map from OT neurons in the four identified areas and examine whether OT projections target specific functional circuits related to distinct behavior control.

Since OT can be released via axons, dendrites, and even neuronal processes (Jurek and Neumann, 2018), we injected a Cre-dependent adeno associated virus 2 (AAV2-CAG-Flex-EGFP) in the four areas of *Ot-Cre* knock-in mice with slightly varying injection sites to cover target areas (N = 15 animals for the PVH, 10 for the SO, 2 for the AN, and 5 for the TU) (Fig. 2A-B). We included male, virgin female, and lactating female mice in our study (see method for more detail), and observed no significant difference between sex or lactating state. Thus, we merged all data from the same anatomical areas. Long-range projection signals from individual injections were then registered onto the Allen CCF and maximum projection data in all samples from each anatomical area were used to represent efferent output for the four areas (Fig. 2C; Movie S2). We found abundant projections from OT neurons in the PVH to the midbrain and hindbrain areas while relatively sparse projection to the diencephalon and telencephalon areas (Fig. 2D)

We then examined whether OT neurons in the four anatomical areas show any distinct projection pattern. Overall, the PVH neurons showed the broadest projection pattern followed by the SO and the AN, which project to a small subset of PVH-OT efferent areas (Fig. 2D-E; Movie S2). The TU-OT neurons did not show any long-range projections. We ask whether OT neurons project to distinct neural circuits related to specific function. Based on known functions of each anatomical region, we found that PVH-OT neurons project to three functional modules that control the internal state, somatic visceral, and cognitive response. Each module contains three circuits. The internal state module contains attention, threat/alert/defense, and sleep/awake circuits (Fig. 2E). The somatic visceral module includes pain, sensory motor, and body physiology/metabolism circuits (Fig. 2E). Lastly, the cognitive control module has learning/memory, reward/value assessment, and reproduction circuits (Fig. 2E). Each circuit is composed of multiple brain regions from the hindbrain, midbrain, thalamus, hypothalamus, cerebral nuclei, and cerebral cortex that process low-to-high order information. For instance, many basal ganglia circuit components including the caudate putamen (CP), globus pallidus (GP), subthalamic nucleus (STN), and substantia nigra (SN) receive OT projection to modulate motor function (see sensory motor regulation in Fig. 2E). PVH-OT neurons project to these areas at varying degrees. Dense projection occurs largely onto the hindbrain (e.g., the dorsal motor nucleus of the vagus nerve; DMX, the parabrachial nucleus; PB), the midbrain (the substantia nigra compacta; SNc), and the hypothalamus (the medial preoptic nucleus; MPN) to directly modulate motor output and sensory input (Fig. 2E). In contrast, the cerebrum (the cerebral nuclei and cerebral cortex) that works as high cognitive controller receives more sparse projection (Fig. 2E). The SO and AN project to a small subset of PVH-OT target areas (Fig. 2E). This data suggests that OT neurons in these two areas can further modulate a subset of the nine functional circuits, albeit less powerfully.

Together, our comprehensive projectome analysis uncovers anatomical substrates to explain pleiotropic effect of OT neurons regulating diverse behavioral outcomes.

### Oxytocin receptor expression showed quantitative mismatch with oxytocin neuronal output

Next, we ask whether expression of a single subtype of the OT receptor (OTR) is quantitively correlated with OT projection target areas to mediate circuit specific OT function. Although most OT projecting areas are known to contain OTR expression (Grinevich et al., 2016), the quantitative relationship between OT projection and OTR expression across the whole brain is currently lacking.

To understand OT-OTR correlation, max OT projectome data from both the PVH and SO were compared to OTR expression in adult mice using a previously validated mouse line, OTR-Venus (Newmaster et al., 2020). A cohort of adult OTR-Venus mice brains were imaged using STPT and mapped OTR expression in the whole adult brain. These mapped OTR positive neurons (magenta in Fig. 3A) were registered onto the same reference brain along with OT-projections (green in Fig. 3A). Overall, the OTR showed high expression in the cortical area with minimal OT projection, while many midbrain and hindbrain regions have strong OT with little OTR expression (Fig. 3A-B; Movie S3). When we examined whether relative projection of OT neurons is correlated with relative OTR density across the entire brain, we found no significant correlation across the whole brain and major brain areas, except for the thalamus and the medulla (Fig. 3C). Overall, our results highlight lack of quantitative and spatial correlation between OT projections and OTR expression in the mouse brain.

Then, how do OTR rich areas (e.g., the isocortex) receive OT signaling without direct OT projection? A previous study suggested that many OTR expressing neurons may receive OT signal non-synaptically via cerebral spinal fluid (CSF) (Zheng et al., 2014). Hence, we examined whether OT projection fibers make physical contact with ventricles. Indeed, we frequently found OT fibers with thick varicosities at the lateral, 3^rd^, and 4^th^ ventricle surface (Fig. 3D). This further suggests that OT signaling may transmit to the brain via the CSF route in addition to direct transmission in target areas.

### OT neurons mainly receive monosynaptic inputs from the thalamus, hypothalamus, and cerebral nuclei

Since OT neurons are known to integrate external stimuli and internal state, we ask whether OT neurons in the PVH and the SO receive mono-synaptic input from sensory and integrative information processing brain areas.

To map brain-wide mono-synaptic inputs in a cell type specific manner, conditional retrograde pseudorabies viruses were injected into the PVH and the SO of the *Ot-Cre* knock-in mice separately (Fig. 4A) (Wickersham et al., 2007). We confirmed the specificity of labeling by performing co-immunolabeling TVA positive neurons with OT and AVP. None of the TVA infected neurons were AVP positive and were largely OT positive (Fig. S2). We also confirmed no leakiness of TVA labeling by injecting TVA in the PVH of adult C57 mice which did not result in any infection (Fig. S2). To confirm G protein dependency for monosynaptic tracing, TVA without G and rabies viruses were injected to the PVH and SO separately and the brains were imaged in STPT (N=1 animal each for the PVH and SO). All the neurons observed were confined to the injection site and both samples were devoid of any long-range input cells. Lastly, we performed another rabies tracing experiment with optimized G and split TVA that are known for improved Cre specificity and tracing (Kim et al., 2016). We found near identical results with this alternative virus approach (Fig. S3) (N=2 animals, each for the PVH and SO). Once we confirmed the validity of our input tracing methods, we used our mapping method to quantify input neurons throughout the whole brain (Kim et al., 2017). To acquire overall inputs to each anatomical area, input signals from multiple independent injections targeting a specific brain region were registered onto the Allen CCF and the max projection of input signals from each anatomical area (N = 6 animals for the PVH and 4 for the SO) were overlaid onto the reference brain (pseudo-colored green for the PVH and magenta for the SO in Fig. 4B-C; Movie S4).

Overall, OT neurons from the PVH mainly receive inputs from the thalamus, hypothalamus, and cerebral nuclei (Fig. 4C). All brain regions providing inputs to the OT neurons also receive output from the OT neurons except the triangular nucleus of septum (TRS), creating reciprocal connections with afferent areas (Fig. 2E and 4D). Noticeably, OT neurons received little input from hindbrain despite strong output to the same area, suggesting that OT neurons provide largely unilateral output to the hindbrain (Fig. 2E and 4D). Moreover, the cerebral cortex provides little to no input to the OT neurons, further supporting very weak direct interaction between cerebral cortical areas and OT neurons (Fig. 4D; Movie S4).

SO-OT neurons receive overall similar input compared to the PVH-OT neurons (Fig. 4D). The broad afferent pattern is in sharp contrast to the very sparse efferent projection of SO-OT neurons to the brain (Fig. 2E and 4D). When monosynaptic input from the PVH- and SO-OT neurons are compared, SO-OT neurons show input from relatively more lateral parts of the brain (Fig. 4C, arrows).

Collectively, we conclude that OT neurons in the PVH and the SO receive similar input from a subset of brain areas that receive majority of input from hypothalamic areas followed by cerebral nuclei and thalamic areas. (Fig. 4D).

### Input-output wiring diagrams of PVH- and SO-OT neurons provide overall neural circuit control patterns

Based on our long-range output and mono-synaptic input data, we constructed input-output circuit diagrams of OT neurons in the PVH and the SO while annotating each brain area based on their functional categories (Fig. 5). PVH-OT neurons project broadly to all nine identified functional circuits throughout the brain, indicating that OT neurons can modulate information processing at different level of circuits with overall stronger influence in the mid- and hindbrain circuits (Fig. 5). In contrast, mid-level circuits including the diencephalon (the thalamus, hypothalamus), the midbrain, and the cerebral nuclei, provide major input to inform action of OT neurons, providing anatomical substrate to perform an integrative role (Fig. 5). SO-OT neurons receive similar mid-level circuit input compared to the PVH-OT neurons while showing limited central projection to the midbrain and pons (Fig. 5). This suggests that SO-OT neurons mainly serve as peripheral hormonal output.

**Figure 5.**
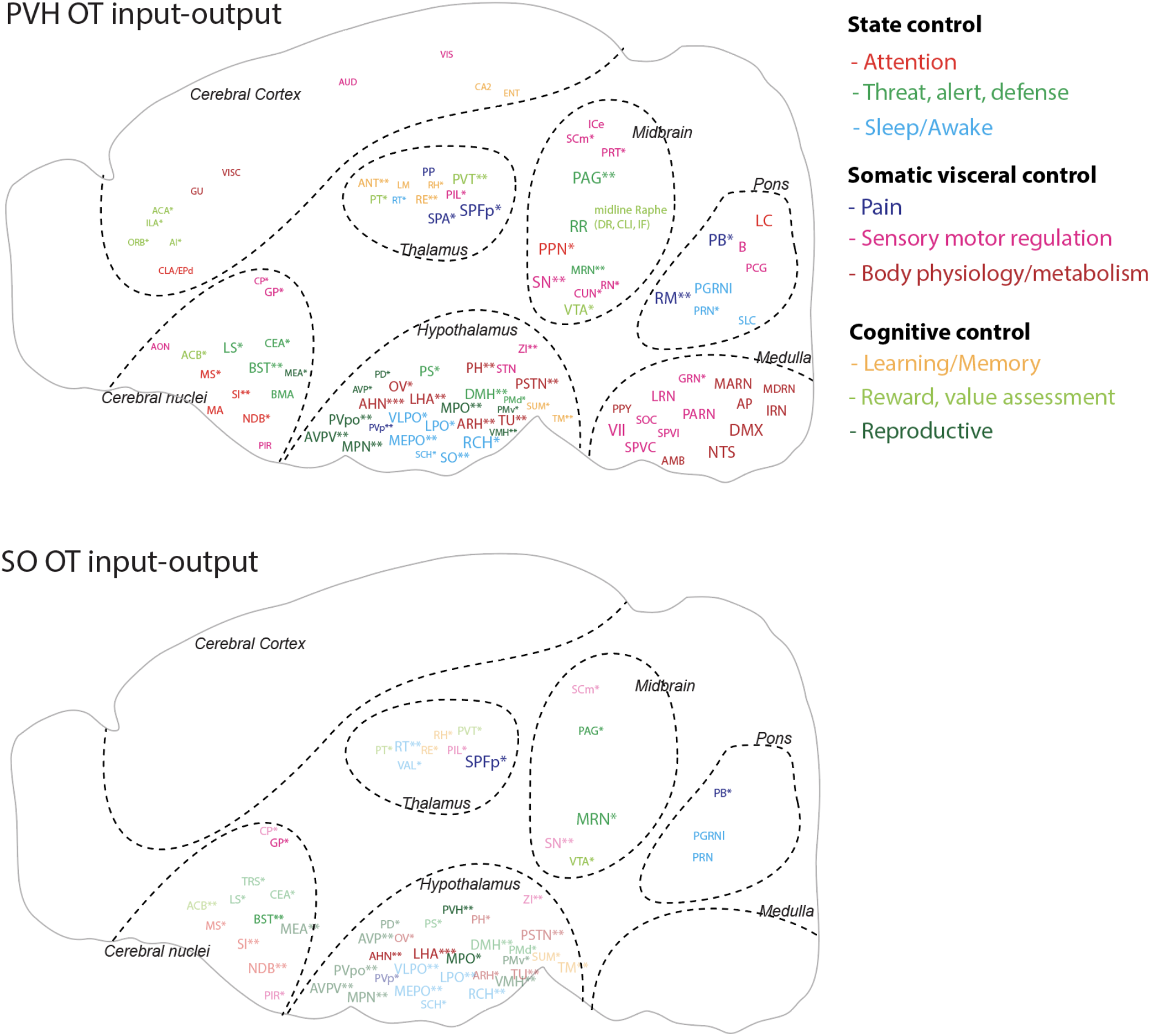
OT input-output wiring diagram. Schematic summary of mono-synaptic input and axonal output connectivity of OT neurons. Color of each ROI is related to nine functional circuits. ROI size is correlated with OT output degree as done in Figure 2. Areas providing the monosynaptic input are highlighted with asterisk (*). Number of * indicated input strength: * < 10 cells, 10 ≤ ** < 100, 100 ≤ ***. Monosynaptic input areas without receiving OT output in the SO-OT map were indicated with semi-transparent fonts. All abbreviations for brain regions can be found in Table 2.

## Discussion

The wiring diagram of the brain is a structural foundation to decipher neural circuits underlying brain function. Here, we present a comprehensive anatomical connectivity map of the hypothalamic OT neurons and their relationship with postsynaptic OTR expression in the whole mouse brain. A quantitative mismatch exists between OT projection and OTR distribution pointing towards abundant non-synaptic OT signaling within the brain. We also identify nine functional circuits with reciprocal or unidirectional connection with OT neurons that serve as anatomical entities to exert varied behavioral control.

OT neurons are mostly located in hypothalamic nuclei with a complex 3D shape (Biag et al., 2012; Madrigal and Jurado, 2021). To examine OT expression intuitively and quantitatively, we devised a 2D flatmap for OT containing hypothalamic regions from an Allen CCF based reference brain while incorporating anatomical labels from the Allen Institute and Franklin-Paxinos (Paxinos and Franklin, 2008; Wang et al., 2020). This approach allows for the interpretation of OT anatomical location from two independently created and commonly used atlases (Chon et al., 2019) and provides an alternative coordinate system to understand anatomical connectivity. In addition to well-described OT neurons in the PVH, SO, and AN, we described another major population in the TU area in the hypothalamus (Jirikowski, 2019). Our 3D immunolabeling independently validated the existence of this extra population. Our anterograde tracing showed that these neurons have almost no central projection, suggesting their contribution to brain information processing is limited. Future studies including ablation studies will help to elucidate the functional significance of this overlooked OT population.

OT signaling is known to modulate many distinct brain functions such as anxiolytic effect, social memory, and attention (Lee et al., 2009; Marlin et al., 2015; Grinevich and Stoop, 2018; Schiavo et al., 2020). By extensively mapping OT efferent processes and clustering brain regions based on known functions, we identified nine functional circuits where OT processes interact to modulate distinct behavioral circuits. Each circuit consists of a set of brain regions processing different behavioral aspects. Thus, our circuit map can help to understand neural entities of OT that modulate different behavioral aspects. Overall, OT circuits provide broad projections to modulate external and internal information throughout the entire brain circuit. For example, we found that OT neurons project to sensory-motor and pain circuits from the hindbrain and midbrain to cerebral cortex and cerebral nuclei. A recent single cell reconstruction study demonstrated that even a single magnocellular OT neuron can make multiple collateral projections to extra-hypothalamic areas to coordinate neuromodulation across functionally related brain circuits (Zhang et al., 2020). These provide anatomical evidence that OT neurons can finely modulate sensory motor processing throughout different circuit levels. Notably, OT neurons project to other neuromodulatory areas such as the locus coeruleus for norepinephrine (alert), the substantia nigra (movement) and the ventral tegmental areas for dopamine (reward), and raphe nuclei for serotonin (emotion), thus serving as a master neuromodulator of neuromodulations (Yoshida et al., 2009; Dölen et al., 2013; Hung et al., 2017; Froemke and Young, 2021). The most well-established role of OT signaling is to promote social behavior (Kemp and Guastella, 2010; Shamay-Tsoory and Abu-Akel, 2015). Social behavior is a complex behavior, requiring coordinated interplay between the sensory system and integrative circuits to generate socially appropriate motor outputs. Extensive connections of OT neurons to somatic visceral, cognitive, and state control modules can help to fine-tune activity of different circuit components to generate enhanced response to socially salient stimuli.

OT gets released through axonal and dendritic projections based on the inputs that OT neurons receive. The presence of large dense core vesicles containing OT at the non-active zones of pre-synapses (Theodosis, 1985; Griffin et al., 2010), absence of evidence for OTR in the postsynaptic membranes, and extremely delayed electrophysiological OT response (milliseconds to seconds) (Knobloch et al., 2012; Knobloch and Grinevich, 2014) collectively support non-synaptic axo-dendritic release of OT. Hence, our projection maps with entire process labeling provide possible release sites of OT throughout the whole brain. We also compared OT total projections (combined data from the PVH- and SO-OT neurons) to OTR expression in the central brain. Although earlier studies mentioned OT-OTR discrepancy, recent studies showed that most OTR expressing areas contain at least sparse OT projection (Knobloch et al., 2012; Grinevich et al., 2016; Mitre et al., 2016; Zhang et al., 2020). Despite a few areas with high levels of both OTR and OT projection (e.g., the paraventricular thalamus), our analysis revealed no significant quantitative correlation between OT and OTR across entire brain regions. For example, the cerebral cortical area contains abundant OTR with little to no OT axons. However, OT can still mediate sensory stimuli in the cortex to modify mouse behavior (Marlin et al., 2015; Schiavo et al., 2020). Previous studies suggest that OTR neurons in the isocortex may receive OT signals indirectly from ventricular pathways via cerebral spinal fluid with delayed and long-lasting effects (Mens et al., 1983; Zheng et al., 2014). Indeed, we found that long-range processes from OT neurons contact the ventricle surface, suggesting potential release of OT into the CSF via long-range processes. Another noteworthy OT-OTR discrepancy is brain regions with abundant OT projection without OTR expression such as sensory related hindbrain areas. Although OTR is a main OT receptor, OT can bind to another receptor to exert its effect. For example, OT can elicit TRPV1 activity in the spinal cord to modulate nociception (Nersesyan et al., 2017). OT modulation in the central nervous system through these non-canonical pathways are under explored and requires further study.

Our OT mono-synaptic input maps showed that the majority of inputs are from the cerebral nuclei, thalamus, hypothalamus, and midbrain with little input from the hindbrain. Particularly, almost all afferent brain regions to PVH-OT neurons also receive efferent projections, suggesting strong reciprocal control of target regions by PVH-OT neurons except the hindbrain for unidirectional output. Abundant bidirectional connections with nine functional circuits suggest that PVH-OT neurons can be an allostatic tool to interactively orchestrate and facilitate social and non-social information processing based on external stimuli and internal state (Quintana and Guastella, 2020). In contrast, despite having similar afferent areas to SO-OT neurons, their limited central projection suggest that SO-OT neurons serve largely as unidirectional hormonal output to the periphery rather than reciprocal circuit modulator.

In summary, our study provides an anatomical foundation to understand diverse functions based on OT neurons in the brain. We deposit all high-resolution imaging data in publicly accessible databases and our website to facilitate data mining. We envision that this OT wiring diagram with quantitative expression data will guide future studies to understand circuit-based mechanisms of OT function and its changes in socially relevant behaviors as well as brain disorders such as autism.

## Acknowledgments

This publication was made possible by an NIH grant R01MH116176 to Y.K. and an NIH grant R24OD18559 to K.C. Its contents are solely the responsibility of the authors and do not necessarily represent the views of the funding agency. We thank Dr. Gloria Choi for kindly sharing OT-Cre transgenic mice and Rebecca Betty for assistance in editing the manuscript. We acknowledge use of computational resources in the High Performance Computing cluster at the Penn State College of Medicine. Fig. 2A and 4A were made by using BioRender.com.

## Contributions

Conceptualization, Y.K.; Data Collection and analysis, S.S., S.M., K.N., M.C., Y.K.; Computer Coding, Y.W.; Web visualization, D.J.V., K.C.; Virus making, T.A.,; Manuscript preparation: S.S., S.M., Y.K with help from the other authors.

