## Supplementary Info for "Whole brain wiring diagram of oxytocin system in adult mice"

Supplementary Information

### Wiring diagram of the oxytocin system in the mouse brain

Son and Manjila et al.,

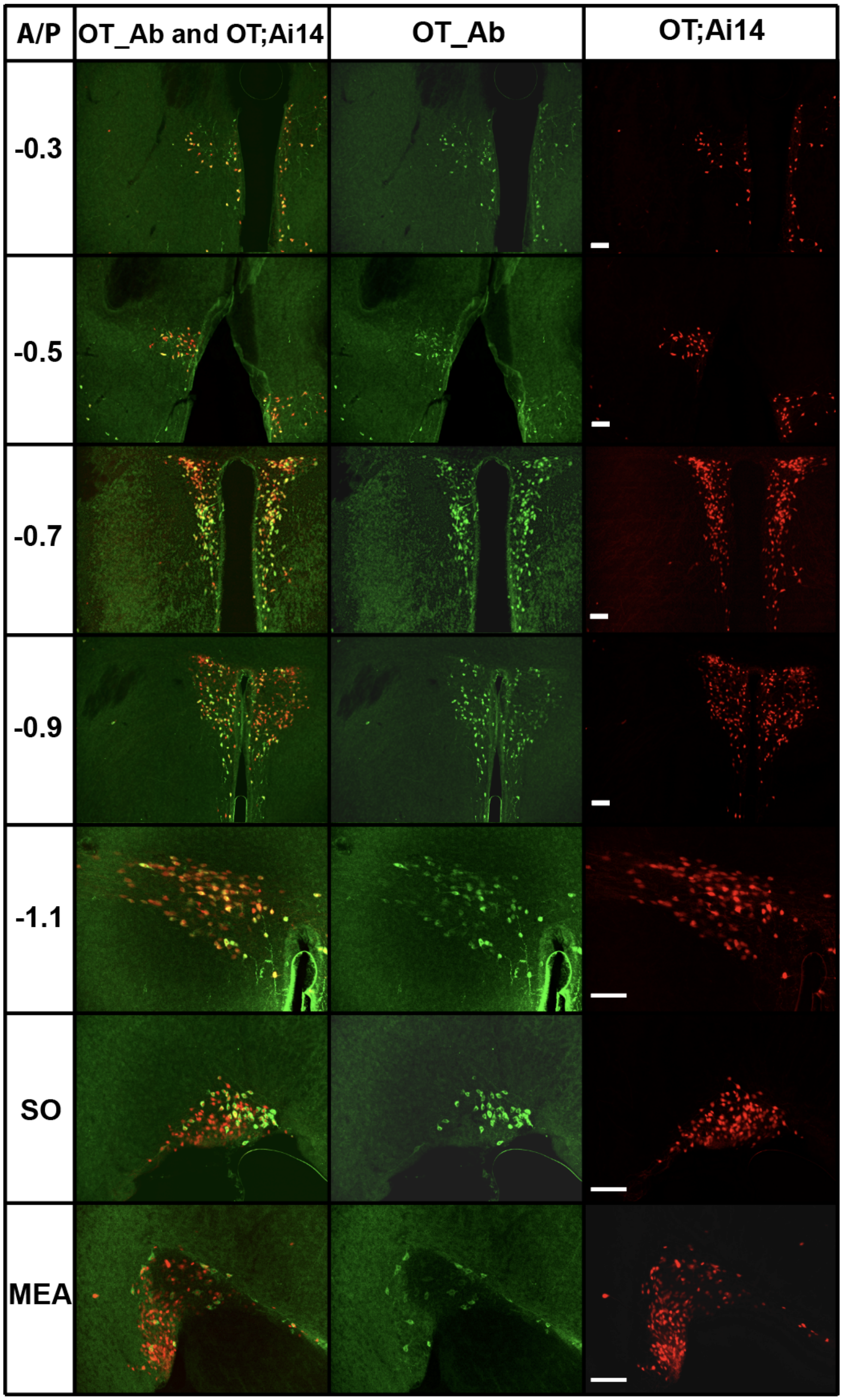


Figure S1. Fluorescent images across 5 levels of the PVH, SO, and MEA. Genetically expressed oxytocin neurons (OT;Ai14) are red and oxytocin immuno staining cellar are labeled with green fluorescent marker are green. Scale bar = 50 µm.

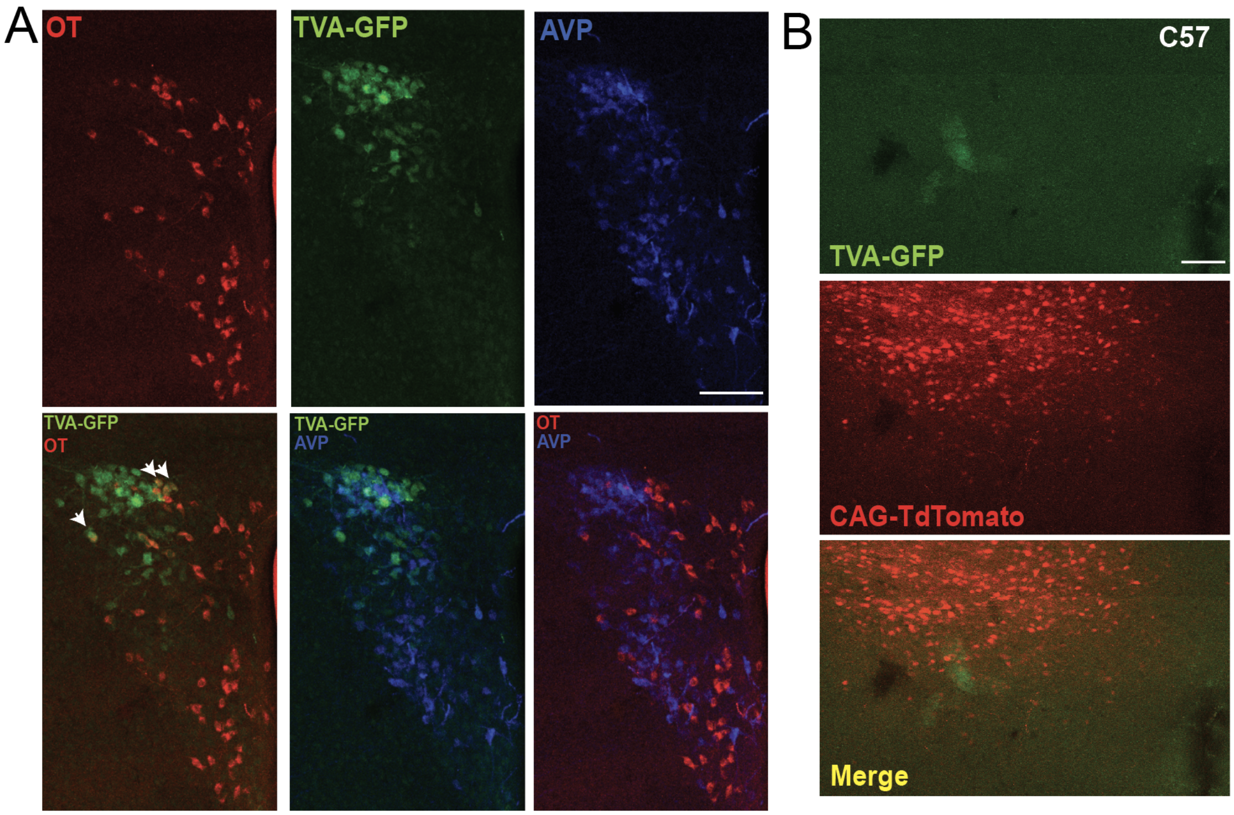


Figure S2. OT and Vasopressin immunolabelling in TVA-GFP injected *Ot-Cre*. (A)TVA-GFP neurons are co-localized with oxytocin (arrows), whereas no vasopressin positive neurons co-localized with TVA-GFP. TVA-GFP labelled in green, OT in red and vasopressin in blue. (B) Specificity of TVA-GFP to infect only Cre positive neurons. TVA-GFP co-injected with CAG-TdTomato in the PVH of C57 mice. TVA-GFP labelled in green and CAG-TdTomato labelled in red. Scale bar = 200 µm.

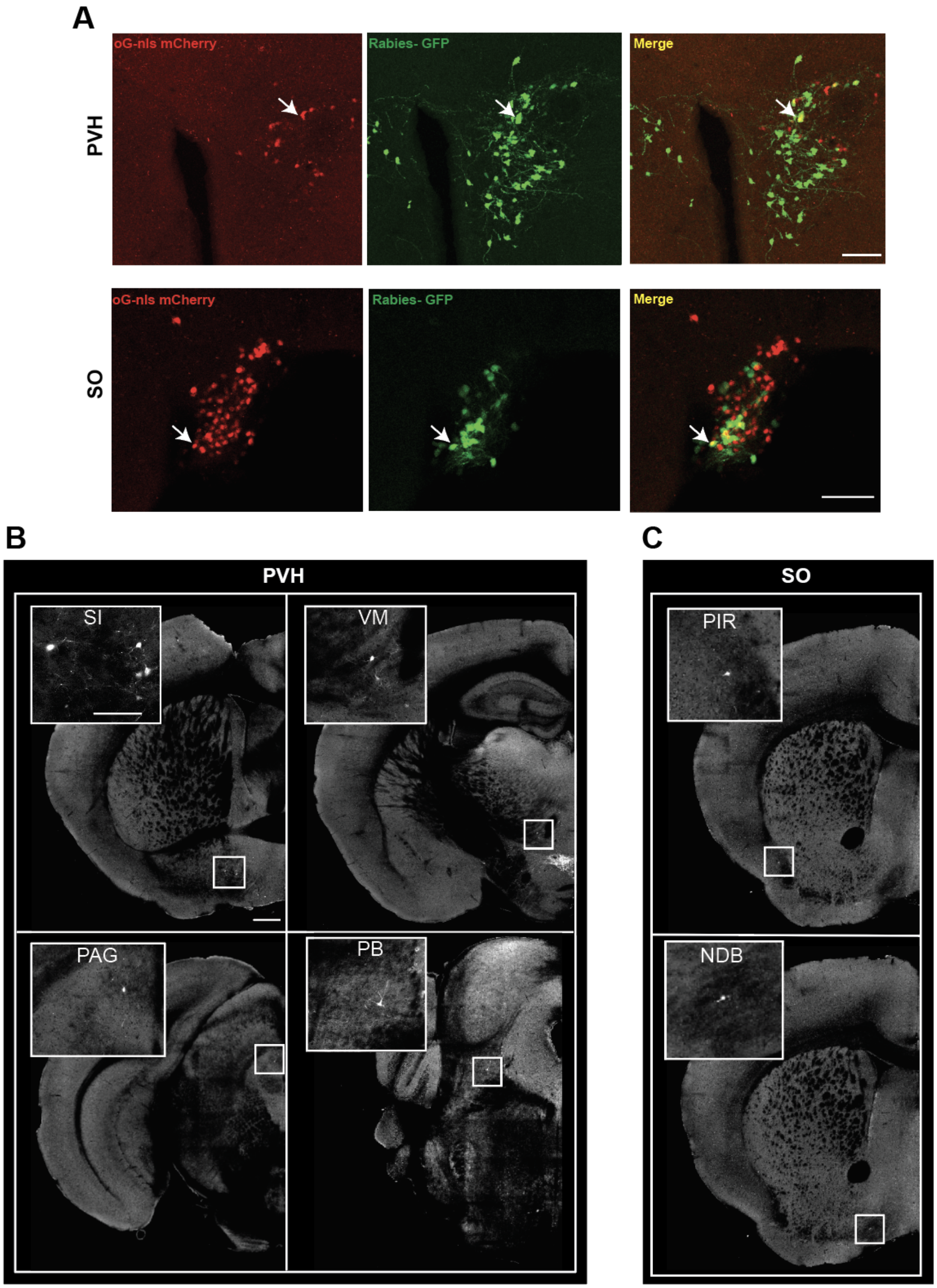


Figure S3. Pseudorabies tracing experiments using split TVA and optimized G injections in the PVH and SO of *Ot-Cre* mice. (A) oG-nls-mCherry marked in red and rabies GFP labelled as green, with overlapping neurons visualized as yellow. (B) High magnification images showing cell bodies of neurons from which PVH and SO OT neurons receive monosynaptic inputs. Scale bar = 200 μm.

**Supplementary Movies**

Movie S1. Oxytocin neuronal expression in the whole brain

Movie S2. Brain-wide projection of oxytocin neurons in the PVH and the SO

Movie S3. Comparison between oxytocin receptor expression and projection of hypothalamic oxytocin neurons.

Movie S4. Monosynaptic input to oxytocin neurons in the PVH and the SO
